# Perivascular and parenchymal fluid characteristics are related to age and cognitive performance across the lifespan

**DOI:** 10.1101/2024.10.14.616343

**Authors:** Kirsten M. Lynch, Rachel M. Custer, Farshid Sepehrband, Arthur W. Toga, Jeiran Choupan

## Abstract

Perivascular spaces (PVS) play a critical role in fluid transfer and waste clearance in the brain, but few studies have explored how alterations to perivascular fluid flow may impact brain maturation and behavior across the lifespan. This study aims to characterize age-related alterations to perivascular and parenchymal fluid flow characteristics across the lifespan in typically developing children (8-21 years) and aging adults (35-90 years) and assess their contribution to cognition. In this study, we employ multi-compartment diffusion models, neurite orientation dispersion and density imaging (NODDI) and tissue tensor imaging (TTI), to quantify free water diffusion characteristics within automatically defined perivascular spaces, the surrounding parenchyma, and at variable distances from the PVS. Our findings show free water diffusion characteristics within the PVS and surrounding parenchyma are associated with age in both developing children and in aging adults. Additionally, age was associated with accelerated change in free water diffusion measures with distance from the PVS. There was no direct effect of free water diffusion measures on cognitive scores across subjects; however, a more complex relationship emerged such that age modified the relationship between free water diffusion measures and cognition. Together, these findings provide evidence of age-associated alterations to fluid flow dynamics and cognition that may be related to the waste clearance system.

## Introduction

The waste clearance pathway is the proposed mechanism through which metabolic waste and toxic substrates are removed from the brain. According to the glymphatic hypothesis introduced by Illif and colleagues (Iliff et al., 2013) in murine models, it is theorized that cerebrospinal fluid (CSF) flows into the brain parenchyma along perivascular spaces, where it mixes with interstitial fluid (ISF) and facilitates the removal of metabolic waste through peri-venous drainage pathways. While much of the direct evidence of the glymphatic hypothesis has been carried out in animal models, recent research in humans show intrathecally-administered tracers undergo CSF-ISF exchange and follow flow patterns consistent with predicted glymphatic mechanisms (Eide and Ringstad, 2024; Ringstad et al., 2018). Therefore, efficient fluid flow is critical for the maintenance of proper brain function and homeostasis.

Enlargement of perivascular spaces (PVS) is considered a marker of impaired waste clearance (Xue et al., 2020), as enlarged spaces can impede fluid transfer from the PVS to the brain parenchyma. Disruptions in fluid flow can lead to the accumulation of toxic substances, including amyloid beta, thus further exacerbating waste clearance dysfunction (Keable et al., 2016). Previous studies have shown increased PVS burden in a variety of neurodegenerative disorders, including Alzheimer’s disease (Banerjee et al., 2017), multiple sclerosis (Kilsdonk et al., 2015) and cerebral small vessel disease (CSVD) (Charidimou et al., 2017, 2013; Doubal et al., 2010; Potter et al., 2015). More recently, studies have also shown increased visibility of PVS in structural MRI over the course of the lifespan, where PVS become progressively wider and more numerous with age (Lynch et al., 2023). However, PVS morphology alone is likely insufficient to fully describe waste clearance efficiency, as several other factors can inform PVS structure (Okar et al., 2023), including underlying vascular size (Zong et al., 2020), microvessel tortuosity (Sun et al., 2024) and *ex vacuo* atrophy and demyelination (Groeschel et al., 2006; Wardlaw et al., 2013). Therefore, methods sensitized to fluid movement in perivascular CSF and parenchymal ISF are needed to more comprehensively describe how fluid flow is altered over the course of healthy aging.

Diffusion-weighted magnetic resonance imaging (DW-MRI) is a non-invasive imaging technique that can provide insight into the molecular movement of water in biological tissue in vivo. DW-MRI is traditionally used to model microstructural features of the brain; however recent studies have shown the diffusion signal is sensitive to fluid flow in the PVS and brain parenchyma (for review, see Barisano et al., 2022). Previous studies in animal models have used diffusion-based MRI sequences to visualize fluid flow along PVS, thus demonstrating the prominent role of vessel pulsatility on fluid movement (Harrison et al., 2018). Additionally, the disruption of fluid transfer between the PVS and parenchyma through inhibition of aquaporin-4 (AQP4) channels are accompanied by alterations in diffusion metrics believed to reflect interstitial fluid stagnation in the brain parenchyma (Debaker et al., 2020; Gomolka et al., 2023). In human studies, researchers have shown altered interstitial fluid behavior in both Alzheimer’s disease (Sepehrband et al., 2019b, 2019c) and CSVD (Jiaerken et al., 2021) in parenchymal tissue, suggestive of waste clearance dysfunction. However, few studies have explored how interstitial fluid dynamics are altered as a consequence of aging and how it may impact cognitive ability in the general population.

Multi-compartment models represent a potentially powerful tool to study fluid flow related to the waste clearance system (Poulain et al., 2023). In the present study, we employ two distinct multi-compartment models, neurite orientation dispersion and density imaging (NODDI) and tissue tensor imaging (TTI), to probe perivascular and interstitial fluid behavior over the course of the lifespan. NODDI is a popular three-compartment model that estimates the intra-cellular, extra-cellular and free water compartments (Zhang et al., 2012). The isotropic volume fraction (fiso) estimates the fraction of a voxel occupied by free, unobstructed diffusion, commonly observed in CSF. TTI is a technique that utilizes a similar approach to free water-eliminated DTI (fwe-DTI), which models the tissue and free water signals as separate tensors (Pasternak et al., 2009). However, unlike fwe-DTI, TTI capitalizes on the observation that diffusion within PVS have directional preference due to the confines of the astrocytic end feet and models the free water tensor as anisotropic (Sepehrband et al., 2019c). Therefore, this technique can distinguish the tissue and free water diffusivities. Together, NODDI and TTI free water measures provide complementary insight into interstitial fluid behavior and can provide important insight into the biophysical processes that give rise to waste clearance alterations in aging.

The goal of this study is to characterize age-related alterations in perivascular and parenchymal fluid flow dynamics using free water diffusion metrics derived from NODDI and TTI over the course of the lifespan. Additionally, we aim to determine whether fluid flow behavior is related to gross cognitive outcomes. We used high-resolution MRI and behavioral data obtained from the Lifespan Human Connectome Projects (HCP) in Development (n=468, 8-21 years) and Aging (n=507, 35-90 years) to separately explore how fluid dynamics are altered in child development and advancing age. We explored fluid flow properties within automatically segmented PVS and the surrounding parenchyma, as well as how interstitial fluid properties in the brain parenchyma were altered with increasing distance from the PVS to quantify PVS-parenchyma fluid exchange. Here, we hypothesize we will observe higher free water content within the vicinity of the PVS and lower free water content in white matter regions furthest from the PVS. Lastly, we explore the behavioral implications of altered fluid dynamics by comparing age-related alterations in perivascular and parenchymal diffusion with global cognitive scores.

## Methods

### Subjects

Data used in this study was acquired through the Lifespan HCP in Development (Somerville et al., 2018) and Aging (Bookheimer et al., 2019). The Lifespan HCP in Development (HCP-D) recruited subjects between 5 and 21 years of age from Boston, Los Angeles, Minneapolis, and St. Louis. The Lifespan HCP in Aging (HCP-A) recruited a large sample of subjects over the age of 36 from multiple locations, including Washington University St Louis, University of Minnesota, Massachusetts General Hospital, and the University of California Los Angeles. To focus on typical development and aging across the lifespan, a liberal threshold for inclusionary criteria was applied. Within HCP-D, subjects were excluded if they had neurodevelopmental disorders or were born premature. Additionally, participants with major psychiatric or neurological disorders, stroke, clinical dementia, severe depression that required extensive treatment, macular degeneration and profound hearing loss were excluded from the study. Subjects with impaired cognitive abilities were excluded from the study according to tiered age-appropriate cut-off scores for cognitive tests (see **Supplementary Data 1** for additional details). Overall, we included 468 subjects (260 F) between 8 and 21 years of age (14.4 ± 3.8) from HCP-D and 507 subjects (298 F) between 35 and 90 years of age (56.4 ± 13.7) from HCP-A in the present study, as previously reported in (Lynch et al., 2023).

### MRI acquisition

At each of the study locations, subjects were scanned on a Siemens 3T Prisma scanner with an 80 mT gradient coil and a Siemens 32-channel Prisma head coil (Harms et al., 2018). Scanning procedures are standardized across all research sites, utilizing identical platforms with E11C software with an electronically distributed protocol. Initial quality assessments found less than 10% of variance was attributed to site differences (Harms et al., 2018). T1w MP-RAGE (voxel size=.8 mm isotropic, 4 echoes per line of k-space, FOV = 256 × 240 × 166 mm, matrix = 320 × 300 × 208 slices, 7.7% slice oversampling, GRAPPA = 2, pixel bandwidth = 744 Hz/pixel, TR = 2500 ms, TI = 1000 ms, TE = 1.8/3.6/5.4/7.2 ms, FA = 8 degrees) and T2w turbo spin echo (TSE) scans (voxel size=.8 mm isotropic, 4 echoes per line of k-space, FOV = 256 × 240 × 166 mm, matrix = 320 × 300 × 208 slices, 7.7% slice oversampling, GRAPPA = 2, pixel bandwidth = 744 Hz/pixel, TR = 3200 ms, TE = 564 ms, turbo factor = 314) were used from each subject to delineate the fine PVS structure. Additionally, multi-shell diffusion weighted images (DWI) with 185 diffusion-encoding directions were acquired across 4 consecutive runs with 28 b=0 s/mm^2^ volumes equally interspersed throughout the scans (multi-band factor = 4, voxel size=1.5 mm isotropic, TR=3.23 s, b=1500 and 3000 s/mm^2^). Scans were acquired with opposite phase-encoding directions (AP and PA) for distortion correction.

### MRI data processing

T1w and T2w structural data were preprocessed with the HCP Preprocessing Pipeline version 4.0.3 (Glasser et al., 2013) using the LONI pipeline version 7.0.3 (Dinov et al., 2009). These steps include gradient distortion correction, rigid body alignment with boundary-based registration (BBR) (Greve and Fischl, 2009) and brain extraction. Freesurfer version 6 (http://surfer.nmr.mgh.harvard.edu/) was run on the T1w images to execute motion correction, intensity normalization and automated Talairach transformation (Reuter et al., 2010; Segonne et al., 2004; Sled et al., 1998). Finally, subcortical white matter was segmented from the T1w volume from the Freesurfer parcellation and used as the brain parenchymal mask for subsequent analyses. dMRI data was preprocessed using the HCP diffusion preprocessing pipeline version 4.1.0 (Sotiropoulos et al., 2013). Data was corrected for motion, eddy-current distortions and susceptibility-induced artifacts using FSL EDDY (version 6.0.3) (Andersson & Sotiropoulos, 2016). The dMRI data was then denoised using a non-local means filter. The b=0 s/mm^2^ volumes were averaged into a single volume and spatially aligned to the T1w and T2w images using a rigid body transformation using ANTs version 2.4.2 with tricubic interpolation (Avants et al., 2008). The resulting registration was then applied to the dMRI metric volumes to ensure spatial correspondence between the PVS masks and microstructural maps.

### Perivascular space segmentation

PVS were visibly enhanced and segmented using an automated image processing approach outlined in (Sepehrband et al., 2019a). First, an enhanced PVS contrast (EPC) was generated from the T1w and T2w images. We performed field bias correction and non-local filtering using image self-similarity properties on the T1w and T2w images. To increase PVS visibility, we then divided the filtered images (T1w/T2w). Next, we segmented the PVS from the EPC using thresholded maps. Briefly, PVS vesselness maps were derived from Frangi filters (default values α = β = 0.5) applied to the EPC (Frangi et al., 1998). The vesselness maps were then thresholded using optimized thresholding (scaled *t* = 1.5) based on the highest concordance and correlation with values derived from expert readers in the HCP Young Adult dataset (Sepehrband et al., 2019a). We have recently found that this thresholding value also performs well in the HCP Lifespan dataset (Lynch et al., 2023). PVS labels with less than 3 contiguous voxels were removed from the total brain mask.

### Diffusion models

Microstructural diffusion maps were computed using the Quantitative Imaging Toolkit (QIT) (Cabeen et al., 2018). Microstructural metrics indicative of free water diffusion behaviors were derived from the multi-compartment models NODDI and TTI. The isotropic volume fraction (FISO) from the NODDI model (Zhang et al., 2012) was calculated within each voxel using the spherical mean technique implemented with QIT (Cabeen et al., 2019). The NODDI model adopts a three-compartment model that distinguishes intracellular, extracellular and CSF compartments. FISO reflects the CSF compartment and represents the fraction of the voxel occupied by fast, isotropic Gaussian diffusion.

TTI is a multi-compartment extension of DTI. While DTI measures reflect the diffusion properties of both tissue and fluid from parenchymal PVS, TTI models separate tensors for the tissue and non-parenchymal fluid (Sepehrband et al., 2019c). TTI measures were first initialized using parameters derived from the bi-tensor model, free water-eliminated DTI (fwe-DTI), using iterative least squares optimization (Hoy et al., 2014). For the TTI procedure, the fluid compartment tensor was required to be axially symmetric with positive diffusivities and modeled as an aligned zeppelin (Sepehrband et al., 2019c). The tissue compartment was re-parameterized with the Cholesky decomposition (Barmpoutis et al., 2009). The free water diffusivity measures derived from TTI (i.e., axial diffusivity (fAD), radial diffusivity (fRD), mean diffusivity (fMD) and fractional anisotropy (fFA)) were extracted from each voxel and used in subsequent analyses (**Figure 1A**).

**Figure 1.**
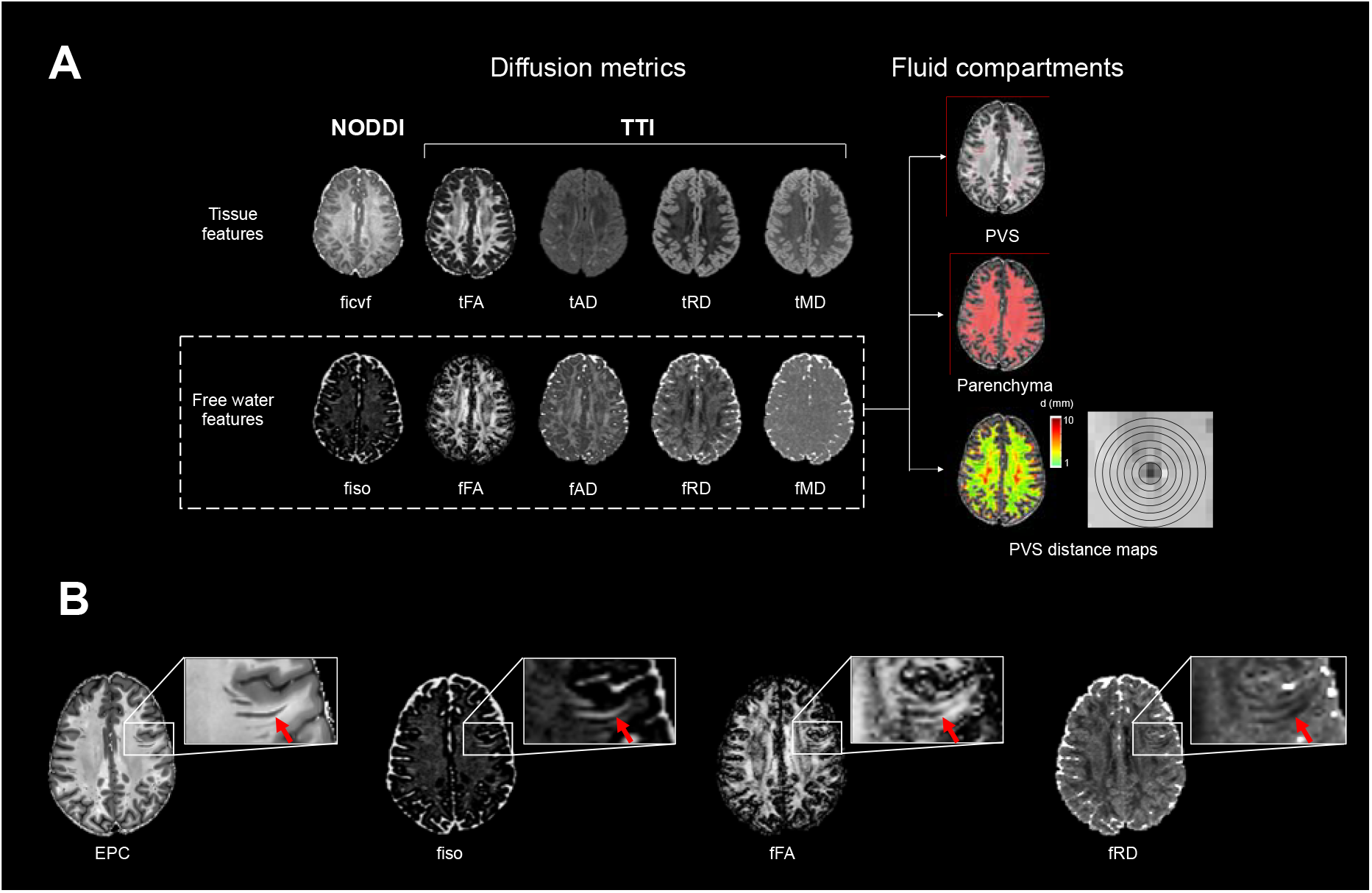
The free water diffusion metric framework used in the present study. (A) The neurite orientation dispersion and density imaging (NODDI) and tissue tensor imaging (TTI) models were run on the diffusion MRI data, yielding the tissue features (ficvf, tFA, tAD, tRD and tMD) and the complementary free water features (fiso, fFA, fAD, fRD and fMD). The free water diffusion features were then averaged within segmented PVS, the PVS-subtracted brain parenchyma, and within 1-mm thick concentric rings surrounding the PVS. (B) The presence of PVS can be observed in an enhanced PVS contrast (EPC) axial slice. These PVS are clearly visible on the same axial slices of fiso, fFA and fRD maps.

The NODDI and TTI measures were averaged within the PVS masks for each subject to obtain estimates of perivascular fluid flow. PVS masks were also subtracted from the white matter masks to obtain parenchymal tissue masks without the contribution of perivascular free water. To avoid partial volume effects of PVS diffusion in the brain parenchyma, we further eroded the parenchymal tissue masks 1 mm around adjacent PVS. Diffusion measures were then averaged for each subject within the parenchyma masks. It is possible that the presence of white matter pathology, such as white matter hyperintensities (WMH), may influence the diffusion metrics, particularly in HCP-A. However, we are unable to segment out WMH due to the absence of FLAIR data in HCP Lifespan. We have previously shown in a subset of HCP-A subjects that 76% of subjects present with no observable WMH, and only 8.5% of subjects presented with either moderate or severe WMH (Lynch et al., 2023). Therefore, we do not expect WMH burden significantly influences the parenchymal diffusion metrics. Lastly, to assess how parenchymal diffusion measures change with distance from the PVS, we derived 10 equidistant rings originating from the PVS masks spaced 1 mm apart and computed diffusion measures within each ring mask (**Figure 1A**). This resulted in separate ring masks for each of the 10 possible distances from the PVS (1mm surrounding PVS, 2mm from PVS, 3mm from PVS, etc.).

### Cognitive scores

All subjects in the HCP-D and HCP-A were administered the NIH Toolbox Cognitive Function Battery (Weintraub et al 2013). We used the total cognitive composite scores, which reflected overall cognition determined from both crystallized (ability to recall past learning abilities) and fluid (ability to apply new learning abilities to novel situations) cognitive abilities.

The NIH Toolbox cognitive assessments consist of 7 test instruments that measure 8 abilities within 6 major cognitive domains. The total cognitive composite score combines performance across cognitive domains to yield a composite score indicative of overall, global cognition. Because the present study focuses on the role of free water diffusion within the entire brain across the lifespan, we hypothesize that alterations to waste clearance function would lead to compromises in global cognitive abilities.

### Statistical analysis

#### Effects of Age on Diffusion Measures within Parenchyma and PVS

Diffusion measures were approximately normally distributed within the parenchymal and perivascular masks (**Supplementary Figure 1**). Bayesian information criteria (BIC) was used to determine the best fit model (linear, quadratic, cubic) to relate the main effect of age on diffusion metrics. Linear models were used to assess the main effect of age on mean parenchymal and perivascular free water diffusion measures, as these models explained the greatest proportion of variance in the data. Sex and scan location were included as covariates of no interest in separate models for HCP-D and HCP-A datasets. The choice to analyze HCP-D and HCP-A separately stems from the fact that we had missing data for participants between the ages of 21 and 35 years of age, alongside the notion that cognitive abilities manifest differently in development versus aging.

#### Effects of Diffusion Measures on Cognition

The role of perivascular and parenchymal free water diffusion processes on cognition throughout the lifespan were assessed using linear models, where we calculated the interaction between diffusion metrics and age, as well as the main effect of diffusion metrics, on total cognition composite scores separately for HCP-D and HCP-A. In order to evaluate both the biophysical free water volume fraction and overall diffusivity of the free water on cognition, we used the metrics fFA and fiso in the analyses. To adjust for multiple comparisons, we employed Bonferroni correction and considered a statistical significance level p<.00625 (2 cohorts * 2 diffusion models * 2 tissue compartments = 8 comparisons). All models include age and sex as covariates.

#### Effect of Distance from PVS on Diffusion Metrics within Parenchyma

To investigate how parenchymal free water varies with distance from the PVS, a linear mixed effects (LME) model was employed using maximum likelihood estimation. This method is appropriate for clustered data, as it treats the multiple measurements of free water at different distances within each subject as correlated observations. The models included a random intercept for each subject, allowing for subject-specific baseline free water measures. The analysis, therefore, examines the average trend of free water change with distance, while accounting for individual variations in baseline free water content. Using LME, we explored the main effect of distance from PVS on diffusion metrics and considered a Bonferroni corrected statistical significance threshold of p=.05/(6 diffusion metrics)=.0083. We also explored the interaction between diffusion metrics (fFA and fiso) and distance from PVS on gross cognition in both HCP-D and HCP-A. To adjust for multiple comparisons, we employed Bonferroni correction and considered a statistical significance level p<.0125 (2 cohorts * 2 diffusion models = 4 comparisons). All models include age and sex as covariates. Years of education was also included as a covariate in models of cognition in analyses of the HCP-A. We did not include this as a covariate in HCP-D analyses as inclusion of both age and years of education in the same model would introduce significant multicollinearity in this cohort, which can lead to unstable and unreliable coefficient estimates. Given the compulsory nature of education in this age range, age itself serves as a strong proxy for educational attainment.

## Results

Free water diffusion measures showed contrast differences between the PVS and parenchyma (**Figure 1B**). Across both cohorts, white matter PVS had significantly higher fiso (*t*(974) = 161.61, *p*<.001), fFA (*t*(974) = 106.42, *p*<.001) and fAD (*t*(974) = 57.45, *p*<.001), and lower fRD (*t*(974) = −−71.56, *p*<.001) and fMD (*t*(974) = −20.67, *p*<.001) compared to the surrounding brain parenchyma (**Figure 2**).

**Figure 2.**
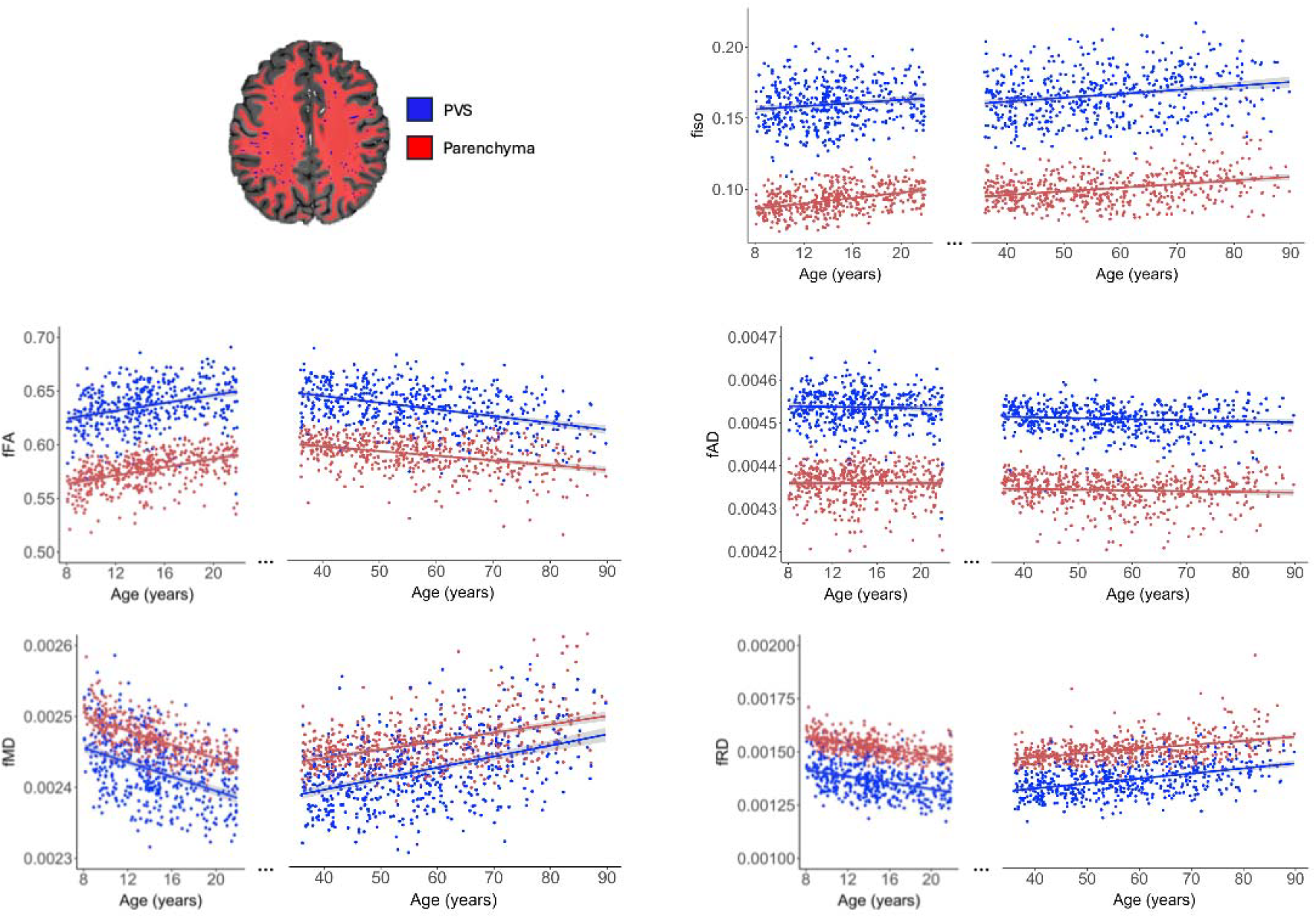
Free water diffusion measures are differentially associated with age across the lifespan. Free water diffusion measures (fiso, fFA, fAD, fMD and fRD) are correlated with age in the developing (8-21 years) and aging (35-90 years) cohorts separately. Individual data points within the PVS (blue) and brain parenchyma (red) are shown, and the best fit line is provided for each tissue compartment.

### Age-related alterations in free water diffusion metrics

Within PVS and the surrounding brain parenchyma, fFA increased with age in development (PVS: *r*=0.29, *p*<.001; parenchyma: *r*=0.43, *p*<.001) and decreased in aging (PVS: *r*=-0.42, *p*<.001; parenchyma: *r*=-0.37, *p*<.001), while fMD and fRD decreased with age in development (fMD – PVS: *r*=-0.29, *p*<.001; parenchyma: *r*=-0.43, *p*<.001; fRD – PVS: *r*=-0.28, *p*<.001; parenchyma: *r*=-0.42, *p*<.001) and increased in aging (fMD – PVS: *r*=0.43, *p*<.001; parenchyma: *r*=0.27, *p*<.001; fRD – PVS: *r*=0.44, *p*<.001; parenchyma: *r*=0.39, *p*<.001) (**Figure 2**). While ficvf showed rapid increases in development and gradual decline in aging (**Supplementary Figure 2**), fiso increased steadily with age across the lifespan (HCD - PVS: *r*=0.13, *p*=.004; parenchyma: *r*=0.39, *p*<.001; HCA - PVS: *r*=0.22, *p*<.001; parenchyma: *r*=0.31, *p*<.001). Because PVS become progressively larger with age (Lynch et al., 2023), we sought to determine whether free water diffusion alterations were due to age-related perivascular dilation. All perivascular free water diffusion measures were significantly associated with the mean diameter of the white matter PVS (**Supplementary Figure 3A**). Within the parenchyma, only fiso was significantly associated with increased PVS diameter (**Supplementary Figure 3B**). The main effect of age on free water tensor metrics remained significant after adjusting for sex and PVS diameter in the model (**Tables 1 and 2**), suggesting that these free water metrics alterations are independent of age-related PVS expansion. Age was not significantly associated with fAD in either development (PVS: *p*=0.41; parenchyma: *p*=.83) or aging (PVS: *p*=.05; parenchyma: *p*=.72).

**Table 1.**
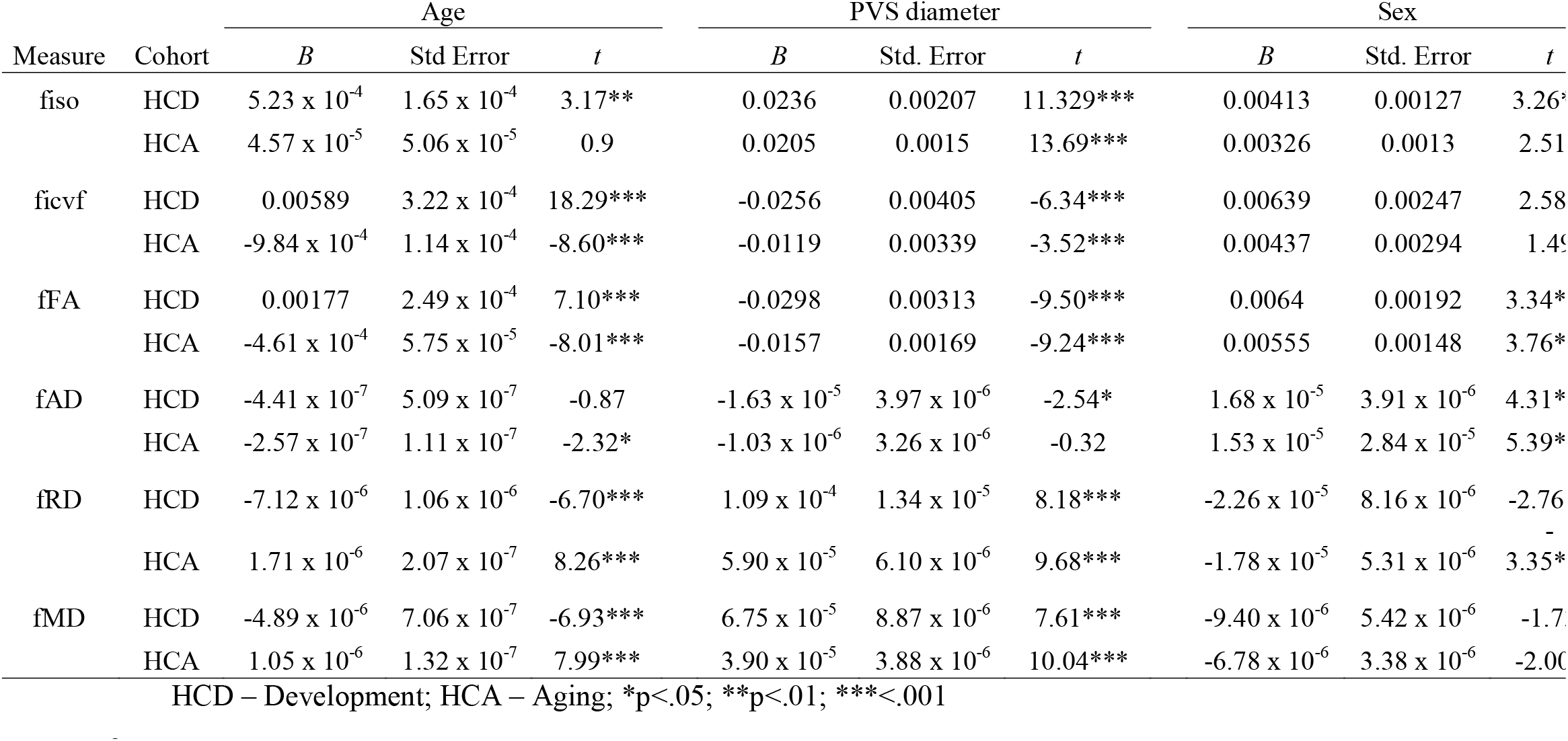
The influence of age, mean PVS diameter and sex on perivascular free water diffusion measures.

**Table 2.** The influence of age, mean PVS diameter and sex on parenchymal free water diffusion measures.

| Measure | Cohort | Age |  |  | PVS diameter |  |  | Sex |  |  |
| --- | --- | --- | --- | --- | --- | --- | --- | --- | --- | --- |
|  |  | *B* | Std Error | *t* | *B* | Std. Error | *t* | *B* | Std. Error | *t* |
| fiso | HCD | 0.00088 | 0.000125 | 6.98*** | 0.00480 | 0.00160 | 2.99** | 0.00270 | 0.000980 | 2.75** |
|  | HCA | 1.73 x 10<sup>-4</sup> | 3.28 x 10<sup>-5</sup> | 5.28*** | 0.00671 | 9.87 x 10<sup>-4</sup> | 6.79*** | 0.00303 | 8.64 x 10<sup>-4</sup> | 3.51*** |
| ficvf | HCD | 0.00598 | 0.00785 | 20.09*** | -0.00799 | 0.00381 | -2.10* | 0.00321 | 0.00233 | 1.38 |
|  | HCA | -6.44 x 10<sup>-4</sup> | 7.78 x 10<sup>-5</sup> | -8.28*** | -0.00210 | 0.00234 | -0.89 | 0.00438 | 0.00205 | 2.14* |
| fFA | HCD | 0.00189 | 0.000187 | 10.14*** | -0.00279 | 0.00239 | -1.17 | 0.00208 | 0.00146 | 1.42 |
|  | HCA | -4.68 x 10<sup>-4</sup> | 5.07 x 10<sup>-5</sup> | -9.23*** | -7.94 x 10<sup>-4</sup> | 0.00152 | 0.52 | 0.00367 | 0.00134 | 2.74** |
| fAD | HCD | -1.33 x 10<sup>-7</sup> | 5.05 x 10<sup>-6</sup> | -0.26 | 7.74 x 10<sup>-6</sup> | 6.45 x 10<sup>-6</sup> | 1.20 | 5.59 x 10<sup>-6</sup> | 3.95 x 10<sup>-6</sup> | 1.41 |
|  | HCA | -1.07 x 10<sup>-7</sup> | 3.83 x 10<sup>-7</sup> | -0.28 | 1.50 x 10<sup>-5</sup> | 1.15 x 10<sup>-5</sup> | 1.31 | 2.11 x 10<sup>-5</sup> | 1.01 x 10<sup>-5</sup> | 2.10* |
| fRD | HCD | -7.05 x 10<sup>-6</sup> | 6.81 x 10<sup>-7</sup> | -10.35*** | 1.38 x 10<sup>-5</sup> | 8.71 x 10<sup>-6</sup> | 1.59 | -8.11 x 10<sup>-6</sup> | 5.33 x 10<sup>-6</sup> | -1.52 |
|  | HCA | 1.89 x 10<sup>-6</sup> | 1.98 x 10<sup>-7</sup> | 9.53*** | 7.96 x 10<sup>-6</sup> | 5.93 x 10<sup>-6</sup> | 1.34 | -9.07 x 10<sup>-6</sup> | 5.21 x 10<sup>-6</sup> | -1.74* |
| fMD | HCD | -4.74 x 10<sup>-6</sup> | 4.37 x 10<sup>-7</sup> | -10.86*** | 1.17 x 10<sup>-5</sup> | 5.58 x 10<sup>-6</sup> | 2.10* | -3.51 x 10<sup>-6</sup> | 3.42 x 10<sup>-6</sup> | -1.03 |
|  | HCA | 1.22 x 10<sup>-6</sup> | 2.13 x 10<sup>-7</sup> | 5.74*** | 1.03 x 10<sup>-5</sup> | 6.38 x 10<sup>-6</sup> | 1.61 | 9.86 x 10<sup>-7</sup> | 5.60 x 10<sup>-6</sup> | 0.18 |

HCD – Development; HCA – Aging; \*p<.05; \*\*p<.01; \*\*\*<.001

**Table 3.** The correlation between age and the PVS/parenchyma ratio of free water diffusion.

| Measure | Cohort | *r* | *p* |
| --- | --- | --- | --- |
| fiso | HCD | -0.348 | <.0001 |
|  | HCA | -0.096 | 0.026 |
| ficvf | HCD | -0.313 | <.0001 |
|  | HCA | -0.444 | <.0001 |
| fFA | HCD | -0.091 | 0.054 |
|  | HCA | -0.167 | 0.0001 |
| fAD | HCD | -0.061 | 0.191 |
|  | HCA | -0.051 | 0.24 |
| fRD | HCD | 0.045 | 0.347 |
|  | HCA | 0.199 | <.0001 |
| fMD | HCD | -0.023 | 0.625 |
|  | HCA | 0.09 | 0.038 |

2 HCD – Development; HCA – Aging

The parenchymal and perivascular diffusion metrics had different age-related slopes. To further investigate this interaction, we explored the relationship between age and the ratio between perivascular and parenchymal fluid characteristics. By using this derived measure, perivascular and parenchymal diffusion trends with age can be more directly compared, as values diverging from 1 indicate increased separation in diffusion metrics between perivascular and parenchymal compartments. The ratio between perivascular and parenchymal fMD, fRD, fFA and fiso was not significantly associated with age (**Supplementary Figure 4**).

### Age-related alterations to parenchymal free water are dependent on distance from PVS

According to the glymphatic hypothesis, fluid enters the brain parenchyma from the PVS. Therefore, we would expect age-related alterations in free water characteristics at variable distances from the PVS if fluid infiltration is altered in aging. We next sought to characterize how free water diffusion measures in the parenchyma are altered with distance from the PVS using masks drawn at sequential 1 mm thick concentric rings around the PVS. Using linear mixed effects models, we found that fFA (*B*=-.014, *t*(8392)=-347.8, *p*<.001), fAD (*B*=-3.4×10^−5^, *t t*(8392)=-355.6, *p*<.001), and fiso (*B*=-.0089, *t t*(8392)=-140.8, *p*<.001) decreased, while fRD (*B*=4.23×10^−5^, *t t*(8392)=141.3, *p*<.001), and fMD (*B*=1.48×10^−5^, *t t*(8392)=243.7, *p*<.001) increased with distance from the PVS, indicative of decreased free water in parenchymal regions furthest from the PVS. A significant interaction between cohort and distance from PVS was observed in all free water metrics, where the slope describing the free water diffusion metrics with distance from the PVS was steeper in the aging cohort compared to the developing cohort (**Figure 3**).

**Figure 3.**
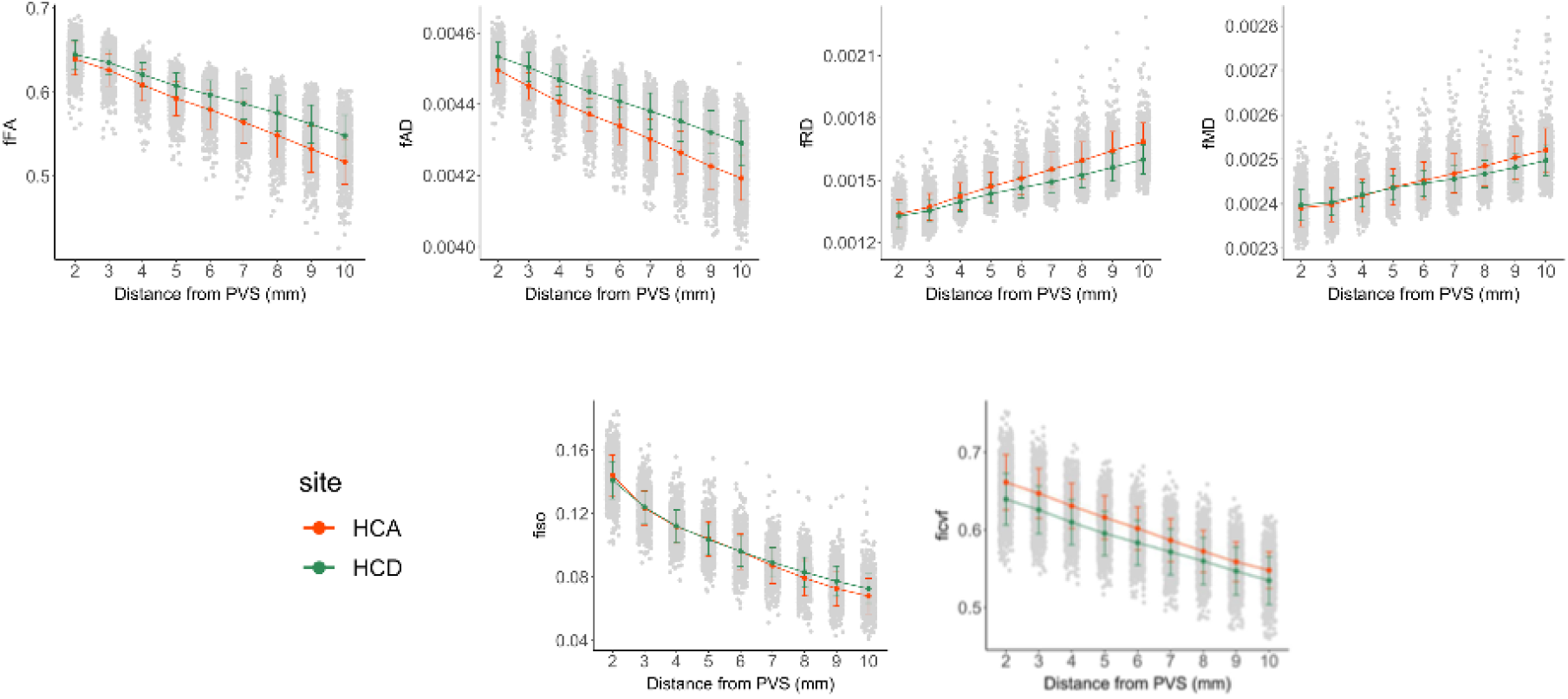
The relationship between free water diffusion measures with distance from PVS differ with age. Free water diffusion measures were averaged within 1-mm concentric circles emanating from PVS and analyzed in HCP-D (green line) and HCP-A (orange line) separately. Significant differences indicate rings around the PVS where diffusion metrics differ between HCP-A and HCP-D. Significant differences in parenchymal fluid characteristics between developing and aging cohorts were observed across diffusion metrics in regions furthest from the PVS (8-10 mm).

### The relationship between free water diffusion measures and cognition is dependent on age

To understand the role perivascular and parenchymal fluid dynamics play in cognitive health across the lifespan, we first tested for the main effect of fFA and fiso on measures of gross cognitive function using the NIH total cognition composite scores using a linear model after accounting for age and sex. Within the aging cohort, no significant associations were observed between cognition and fFA in the PVS (*t*(503)=.442, *p*=.66) and surrounding parenchyma (*t*(503)=1.23, *p*=.22). We then tested for the interaction between diffusion metrics (fFA and fiso) and age on gross cognitive function after controlling for main effects and sex. In the developing cohort, a significant interaction was observed between total cognition and fFA in the PVS (*t*(464)=-4.09, *p*=5.0 × 10^−5^) and parenchyma (*t*(464)=-3.94, *p*=9.9 × 10^−5^), such that the slope is decreased in older adolescents compared to younger children (**Figure 4A and B**). To quantify the strength of the interaction between fFA and age, we calculated the semi-partial correlation (sr) between the interaction term and cognitive performance (PVS: *sr*=-0.14; parenchyma: *sr*=-0.15). We found no significant association between total cognition and perivascular and parenchymal fiso in either cohort after controlling for the effects of age and sex (**Table 4; Supplementary Figure 5A and B)**.

**Table 4.** The interaction between age and diffusion metrics on total cognition composite scores as assessed with the NIH Toolbox.

| Cohort | Region | Metric | metric |  | age |  | metric*age |  |
| --- | --- | --- | --- | --- | --- | --- | --- | --- |
|  |  |  | F | p | F | p | F | p |
| HCD | PVS | fFA | **13.19** | **0.0003** | **20.24** | **<.0001** | **15.54** | **<.0001** |
|  |  | fiso | 0.79 | 0.37 | 0.71 | 0.4 | 0.64 | 0.42 |
|  | Parenchyma | fFA | **12.07** | **0.0006** | **20.92** | **<.0001** | **16.7** | **<.0001** |
|  |  | fiso | 1.42 | 0.23 | **11.82** | **0.0006** | 2.77 | 0.09 |
| HCA | PVS | fFA | 0.2 | 0.66 | 0.06 | 0.81 | 0.01 | 0.94 |
|  |  | fiso | 0.44 | 0.51 | 1.26 | 0.26 | 0.26 | 0.61 |
|  | Parenchyma | fFA | 1.52 | 0.22 | 0.25 | 0.61 | 0.42 | 0.52 |
|  |  | fiso | 0.53 | 0.47 | 0.74 | 0.39 | 0.09 | 0.76 |

HCD – Development; HCA – Aging

**Figure 4.**
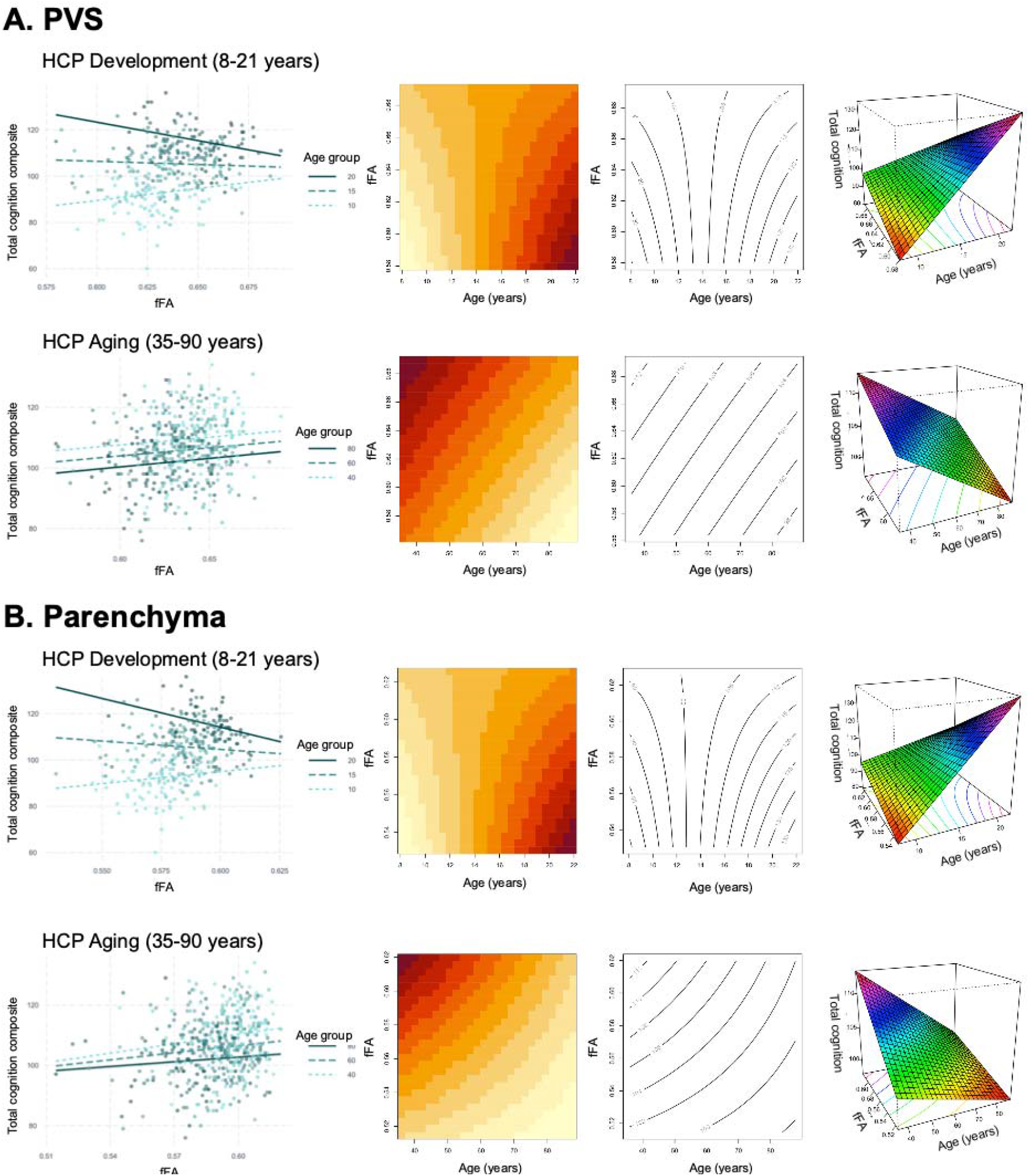
The relationship between free water diffusion and cognition differs across ages. The relationship between fFA and the NIH total cognition composite score (Total Cog) is shown for the (A) PVS and (B) parenchyma. A significant interaction between fFA and age is observed in HCP-D, but not HCP-A. (Left) Individual subjects are shown as points color-coded by age, and representative regression lines are shown for the youngest, middle and oldest ages (from light to dark) for each cohort. Visualization of the three-dimensional nature of the continuous-by-continuous interactions using (middle) matrix plots with contour lines and (right) perspective plots. The contour lines of the matric plots describe the joint distribution of fFA and age, where the area of each binned rectangle in the plot is proportional the frequency of the corresponding combination of variables. The curved, non-parallel nature of the contour lines in the interaction plots for HCP-D (Top A, B) indicates an interaction is observed, while the parallel contour lines for HCP-Aging (Bottom A, B) demonstrates a simple additive relationship. The perspective plots provides a 3D surface-based representation where the height of the surface at any point (x,y = age, fFA) represents the predicted value of the outcome variable (z = cognition). The curved surface observed in HCP-Development (Top A, B) indicates an interaction, while the planar surface observed in HCP-Aging (Bottom A, B) indicates a simple additive relationship.

Next, we assessed whether diffusivity variations with distance from PVS inform cognition across the lifespan using linear mixed effects models. No significant interaction between cognition and distance from the PVS was observed for either fiso (*t*(2576)=1.86, *p*=.06) or fFA (*t*(2576)=-1.17, *p*=.24) in the developing cohort (**Figure 5A**). In the aging cohort, we found a significant interaction between distance from PVS and total cognition scores on fiso (*t*(3793)=5.88, p=4.5 × 10^−9^, sr=.078) and fFA (*t*(3793)=6.56, *p*=5.9 × 10^−11^, *sr*=.019) after controlling for age and sex (**Figure 5B**), and these results remained significant after accounting for years of education in the model for both fiso (*t*(3793)=5.83, *p*=5.9 × 10^−9^, *sr*=.053) and fFA (*t*(3793)=6.53, *p*=7.5 × 10^−11^, *sr*=.019). For fiso and fFA in the aging cohort, free water diffusion measures had steeper decreases with distance from PVS in subjects with lower cognitive performance (**Figure 5**).

**Figure 5.**
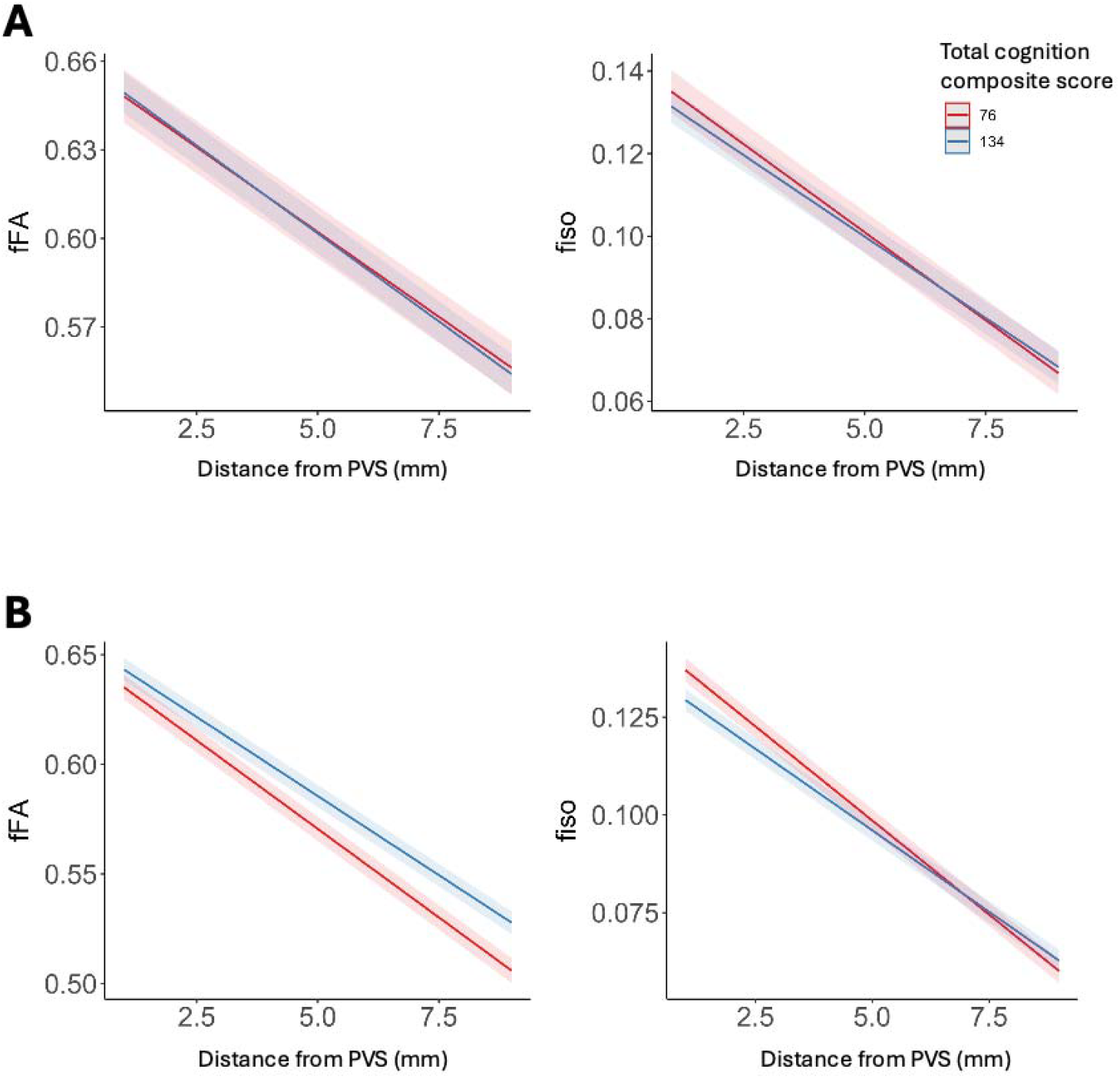
Free water diffusion measures undergo greater alterations with distance from PVS in subjects with lower cognitive performance. The interaction between NIH Toolbox total cognition composite scores and distance from PVS on parenchymal free water diffusion measures (fFA and fiso) are shown for (A) developing children (8-21 years) and (B) aging adults (35-90 years). The estimated regression lines are derived from linear mixed effects models that account for age and sex in the model. Estimated regression lines are shown for 1 standard deviation below (score of 76) and 1 standard deviation above (score of 136) the mean (score of 100).

## Discussion

In the present study, we used free water diffusion MRI metrics derived from multi-compartment models to assess age-related alterations in perivascular CSF and parenchymal interstitial fluid flow across the lifespan. We found age-related alterations in free water diffusion measures that are distinct from the typical microstructural lifespan trajectories. Additionally, we found age-related differences in parenchymal interstitial fluid content with distance from the PVS, where much sharper drop-offs in fluid content are observed in aging adults compared to development. Lastly, fluid behavior in the parenchyma and PVS differentially predicted cognitive performance across age ranges.

Our findings reiterated previous studies that show perivascular and parenchymal fluid compartments can be visually and statistically separated using free water diffusion maps (Jiaerken et al., 2021; Sepehrband et al., 2019c). Our findings show the free water volume fraction is highest in the PVS, presumably due to the increased content of freely diffusing fluid within the PVS compared to the hindered barriers of the brain parenchyma, replete with axons, glia and metabolic debri in the extracellular space. Additionally, we found free water anisotropy is highest in the PVS compared to the parenchyma. This is likely attributed to the preferential movement of unhindered fluid along the length of the PVS compared to the more tortuous path ISF must take through the brain parenchyma. Previous studies have indeed demonstrated that perivascular fluid systematically biases the traditional DTI signal due to the presence of fast diffusion of CSF within the confines of the PVS (Sepehrband et al., 2019c).

We found perivascular and parenchymal fiso increase with age from childhood through advancing age. The fiso lifespan trajectory also appears to be uncoupled from ficvf, echoing previous studies of white matter microstructural alterations over the course of the lifespan. These previous studies showed early and rapid increases in parenchymal neurite density in development (Lynch et al., 2020), followed by a slower decline with advancing age (Beck et al., 2021; Nazeri et al., 2015). It would be reasonable to assume that the free water volume fraction may be inversely correlated with ficvf, as increased myelination in development would reduce the extracellular space while neuronal atrophy in aging would increase the extracellular space. However, our findings demonstrating that the fiso lifespan trajectory in the parenchyma is uncoupled from the ficvf trajectory. This may reflect a more distinct process indicative of the important glymphatic process of bulk fluid flow, which refers to the coordinated, directional movement of CSF and ISF through the PVS and brain tissue (Bohr et al., 2022). These findings are also independent of PVS diameter. Therefore, increased bulk fluid flow within and around PVS is not merely a by-product of the morphological enlargement of the spaces that contain freely diffusing fluid (Lynch et al., 2023). We also found fluid anisotropy increases in childhood and decreases in advancing age, due predominantly to fluctuations in the radial (fRD) and mean (fMD) diffusivities. Together, these findings suggest childhood and adolescence are characterized by increased free water content (fiso) with increased directional preference of free water diffusion (fFA, fRD and fMD), while aging is characterized by increased free water content (fiso) with decreased directional preference of free water diffusion (fFA, fRD, fMD). These diverging findings point towards different free water behaviors in development and aging. Increased free water anisotropy in development may indicate the healthy functioning of CSF/ISF perfusion into parenchymal tissue to aid in the removal of metabolic waste. Meanwhile, increased free water content with decreased free water anisotropy in older adults may indicate increasing perivascular and parenchymal fluid stasis due to age-related impairment to CSF flow dynamics in the waste clearance system (Jiang-Xie et al., 2025). Interestingly, these alterations are accompanied by similar trends in the tissue tensor. Therefore, it is possible that the estimated fluid tensor is influenced by the microscopic boundaries of the surrounding parenchymal tissue. The similar age-related trends observed in free water diffusion metrics in the parenchyma and PVS raise the possibility of shared underlying processes affecting fluid dynamics in these compartments. Passive fluid movement from the PVS to parenchyma is likely driven by a combination of hydrostatic pressure differences (Benveniste et al., 2017) and pulsatile flow (Klarica et al., 2019), and as such, the balance of fluid content between these compartments may be maintained over time in healthy subjects.

To better understand how perivascular and parenchymal fluid contributes to the free water signal in development and aging, we explored the spatial distribution of free water values with respect to the PVS. We observed a reduction in free water anisotropy, as shown by reduced fFA and fAD and increased fRD and fMD, and volume, as shown by reduced fiso, in the brain parenchyma with distance from the PVS. Our findings suggest the amount and preferred direction of bulk flow is lowest in regions where enlarged PVS are not detected. Our findings are supported by a recent study in patients with cerebral small vessel disease (CSVD) and older subjects, where they observed a similar pattern of reduced fiso with distance from the PVS (Jiaerken et al., 2021). We found the difference in free water tensor metrics from the PVS to the periphery is greater in older subjects compared to developing children and adolescents, which may correspond to less stable fluid homeostasis in aging through reduced arterial pulsatility and disrupted blood-brain barrier permeability (Iliff et al., 2013). Together, this would increase resistance to fresh CSF delivery into the parenchyma, and result in the build-up of metabolic waste and fluid stasis, leading to reduced directional preference.

To determine if age-related alterations in free water diffusion measures have functional consequences, we explored the interplay between age, diffusion metrics and gross cognitive performance. We observed an age-dependent relationship between fFA and cognitive performance in the developing cohort, but not the aging cohort, where the correlation between fFA and cognitive performance in both the parenchyma and PVS becomes increasingly negative with age. While it seems counterintuitive that the positive relationship between fFA and cognition would start to weaken in older adolescence, even before the decline in aging sets in, the relative importance of efficient waste clearance for various cognitive domains might shift as cognitive abilities undergo significant changes over the course of adolescence and there may be periods where cognitive gains are less directly tied to further increases in fFA. Additionally, it is possible that fFA may be capturing a combination of factors. Adolescence is a period of intense synaptic pruning and refinement and the removal of unnecessary axons may change the underlying geometric conformation of the brain, thus contributing to a more optimal, but reduced fFA with increasing cognition. Future studies should further explore the contribution of PVS structure and waste clearance function on the developing brain and should differentiate between different cognitive domains, as gross cognition may obscure potentially relevant associations with free water metrics.

(Hamilton et al., 2021; Tang et al., 2022)(Nguyen et al., 2014; Raitamaa et al., 2024)(Santisteban et al., 2023)In aging adults, a more subtle relationship between free water diffusion measures and gross cognition was observed, wherein we found a distance-dependent relationship between cognitive performance and the diffusion metrics fiso and fFA, where subjects with worse cognitive performance had a steeper rate of decrease in diffusion metrics in the parenchyma with distance from the PVS. Therefore, our results suggest free water diffusion measures exert a more subtle and localized influence on cognitive performance in aging subjects. The relationship between the structural components of the waste clearance system and cognition have been previously explored; however, evidence is mixed. Previous studies have shown the number of MR-visible PVS is associated with worse performance in global cognitive function (Libecap et al., 2022), information processing and executive function (Passiak et al., 2019). However, in a large meta-analysis, it was found that PVS counts were not associated with cognitive deficits in the general population (Hilal et al., 2018). However, this is the first study, to our knowledge, that interrogates this question using diffusion MRI measures of bulk fluid flow in both children and aging adults.

Our study explores the influence of diffusion MRI-derived measures of fluid flow properties in the PVS and parenchyma using large population-based datasets that cover the lifespan from development through advancing age. However, several limitations should be addressed. First, the present study is cross-sectional, and future studies should employ longitudinal designs to establish the relationship between fluid flow properties and changes in cognition. Additionally, while others have demonstrated the utility of using the NODDI model to probe the free water diffusion compartment as a measure of fluid flow properties (Jiaerken et al., 2021), additional models derived from ultra-low gradient strength diffusion MRI acquisitions, such as intra-voxel incoherent motion (IVIM), may provide improved sensitivity for bulk flow in the brain parenchyma (Wong et al., 2020, 2017). Since the maximum gradient strength used in the present study may reflect the lower bound of non-Gaussian diffusion (Hoy et al., 2014), it is possible that the bitensor TTI model may be sensitive to some non-Gaussian behaviors such as that observed in restricted diffusion compartments. Together, further studies are needed to better understand the robustness of free water diffusion techniques in measuring fluid flow related to waste clearance using clinically available techniques. Our study also does not contain data for subjects between the ages of 21-35 years. Previous studies using different imaging modalities and metrics have determined that developmental peaks for white matter properties are often observed during this age range (Bethlehem et al., 2022). Therefore, our results may be missing important developmental dynamics during this period.

We demonstrate age-related alterations in free water diffusion metrics indicative of fluid flow properties in the PVS and surrounding parenchyma over the course of the lifespan. The results from this study suggest age-related PVS enlargement is accompanied by alterations in fluid exchange between the PVS and parenchyma and may indicate changing waste clearance functionality over the course of the life. Our findings also show perivascular and parenchymal fluid characteristics are predictive of cognitive outcomes during child development, where increased free water may lead to preferential impairments to cognition in older adolescents. Our findings therefore suggest altered fluid characteristics in the maturing brain may disrupt cognitive development, potentially due to inefficient waste clearance.

## Data and Code Availability

All data used in this study is available through the Human Connectome Project (HCP) (https://www.humanconnectome.org/). The Lifespan HCP datasets (HCP Aging and HCP Development) can be accessed through the Connectome Coordination Facility (CCF) in the NIMH Data Archive (NDA) at https://nda.nih.gov/ccf (DOI: 10.15154/1524651).

## Author Contributions

<u>Kirsten M. Lynch</u>: conceptualization, methodology, formal analysis, investigation, writing – original draft, writing – review & editing, visualization; <u>Rachel M. Custer</u>: formal analysis, data curation, writing – review & editing; <u>Farshid Sepehrband</u>: methodology, software, resources, writing – review & editing; <u>Arthur W. Toga</u>: supervision, project administration, funding acquisition; <u>Jeiran Choupan</u>: software, writing – review & editing, supervision, project administration, funding acquisition.

