## Supplementary for "Perivascular and parenchymal fluid characteristics are related to age and cognitive performance across the lifespan"

Supplementary Data

**Supplementary Data 1 – Cognitive Cut-off Scores**

When recruiting participants aged 60 and above, the HCP Aging study implemented a cognitive screening process to ensure participant competence. They utilize the modified Telephone Interview for Cognitive Status (TICS-M), a validated cognitive assessment tool previously described by de Jager and colleagues (de Jager et al 2003). Eligibility was determined by achieving a minimum score of 30, with score adjustments accounting for educational background. Specifically, the scoring mechanism provides additional points for lower educational levels (5 points for less than 8 years of schooling, 2 points for 8-10 years of schooling) and subtracts points for higher education levels (2 points deducted for 16 or more years of schooling). For participants over 80 years old, where standard normative data may be limited, they employed an alternative screening approach. Candidates must successfully complete critical orientation questions, and those who do not meet the standard score threshold but pass these critical items undergo an additional capacity to consent assessment.

During the inclusion and exclusion assessment, the Montreal Cognitive Assessment (MoCA) (Nasreddine et al., 2005) was administered and inclusion is based off the determined threshold for their age group (36-79 years: 19; 80-89 years: 17; >90 years: 16). Inclusion of participants with below-threshold scores were permitted in subjects older than 80 years of age with macular degeneration and hearing loss if able to communicate via microphone in the scanner to include individuals without mild cognitive impairment or dementia that may have low scores.

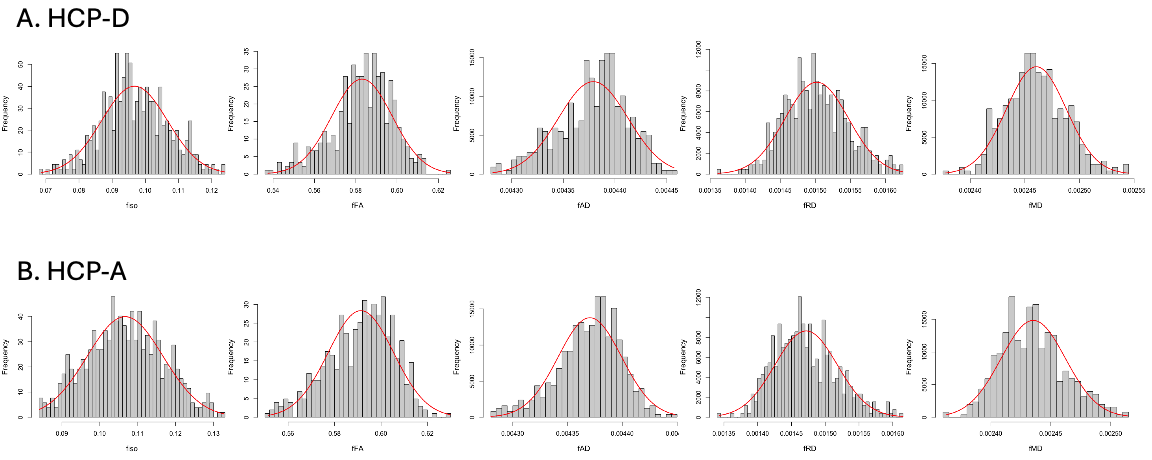

**Supplementary Figure 1.** Distribution of free water diffusion values in (A) HCP-D and (B) HCP-A. The normal curve is overlaid on the histogram in red.

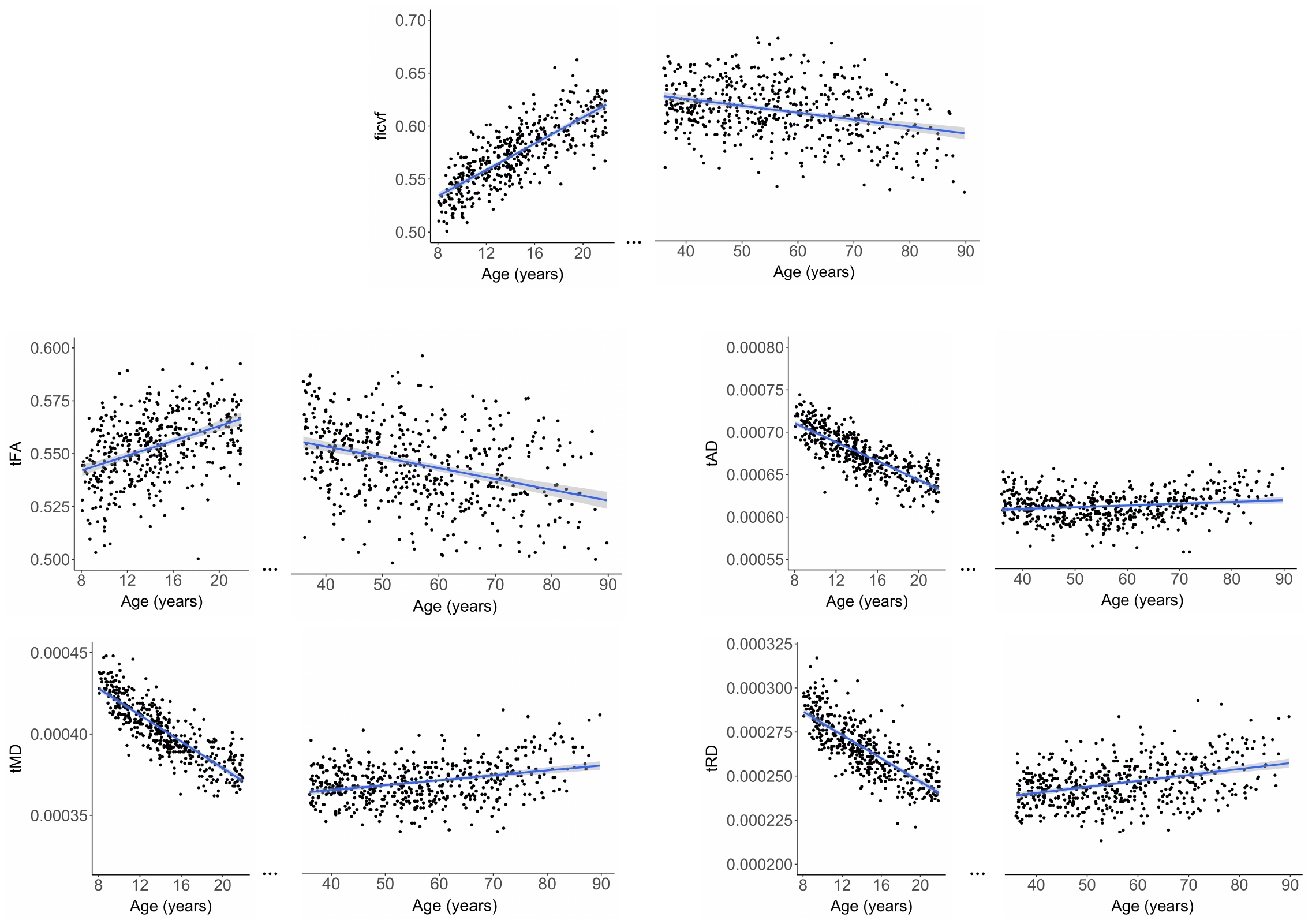

**Supplementary Figure 2**. The main effect of age on microstructural tissue measures derived from NODDI (intracellular volume fraction – ficvf) and TTI (tissue tensor metrics tFA, tAD, tRD and tMD). Regression lines are drawn separately for the developing (8-21 years) and aging cohorts (35-90 years).

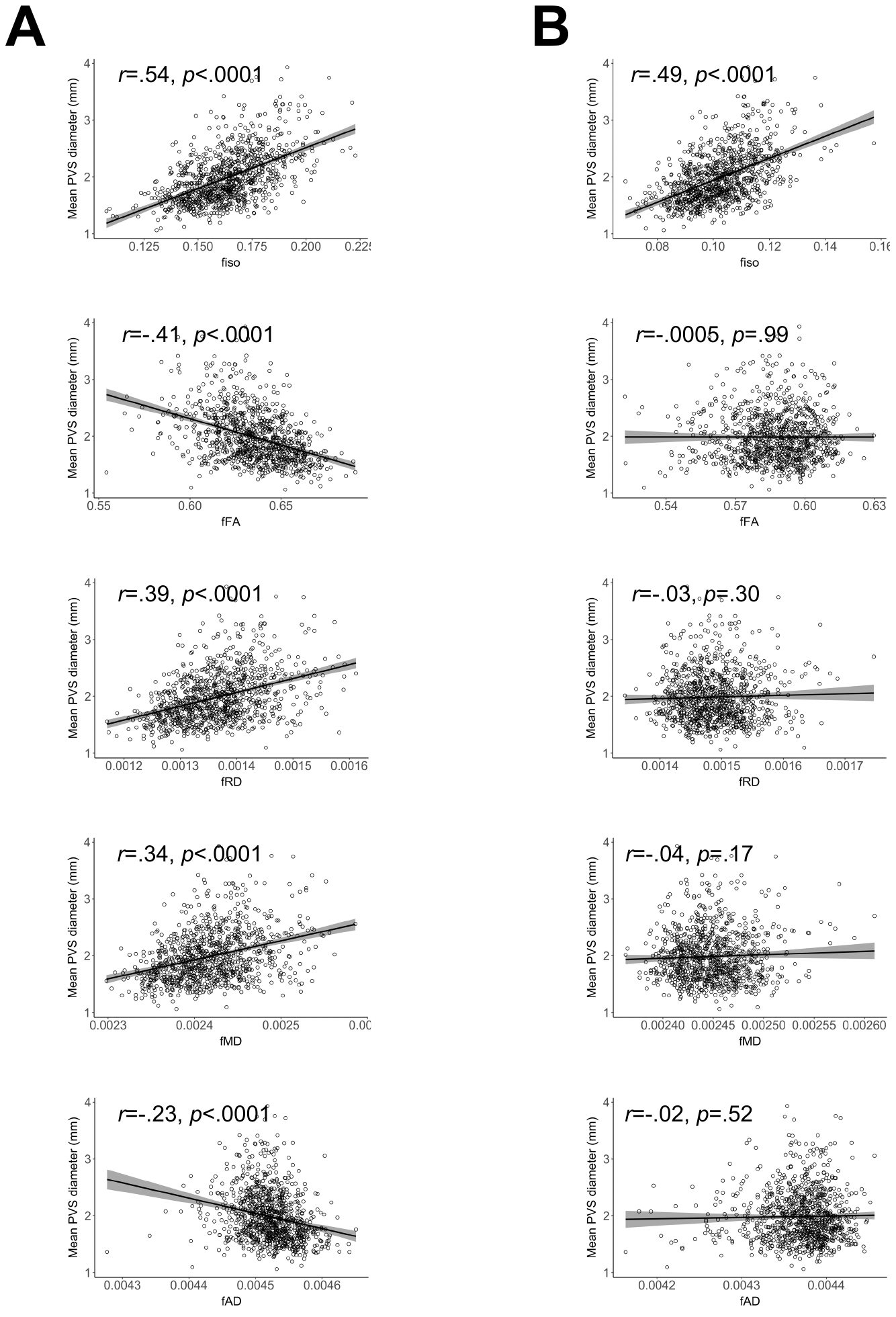

**Supplementary Figure 3**. The association between mean PVS diameter and free water diffusion measures in the (A) PVS and (B) parenchyma. The Spearman’s correlation coefficient (*r*) and *p*-value are shown for each metric.

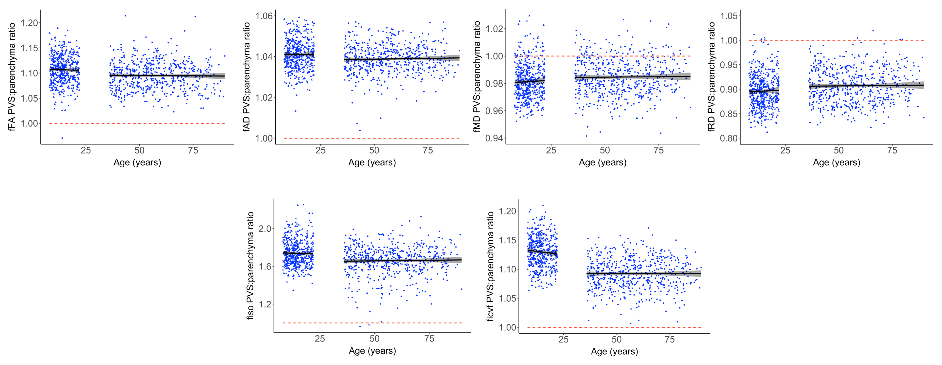

**Supplementary Figure 4.** The relationship between age and the ratio between PVS and parenchymal free diffusion. Free water diffusion measures expressed as the ratio between PVS and parenchyma are correlated with age. A value of 1 denotes subjects where the diffusion metric is the same between the PVS and parenchyma (dashed red line). Values greater than one indicate metrics that are higher in the PVS compared to the parenchyma, and values lower than one indicate metrics that are lower in the PVS compared to the parenchyma. Significant correlations with age are indicated where the correlation coefficient (*r*) is shown. * *p*<.05, ** *p*<.01, *** *p*<.001.

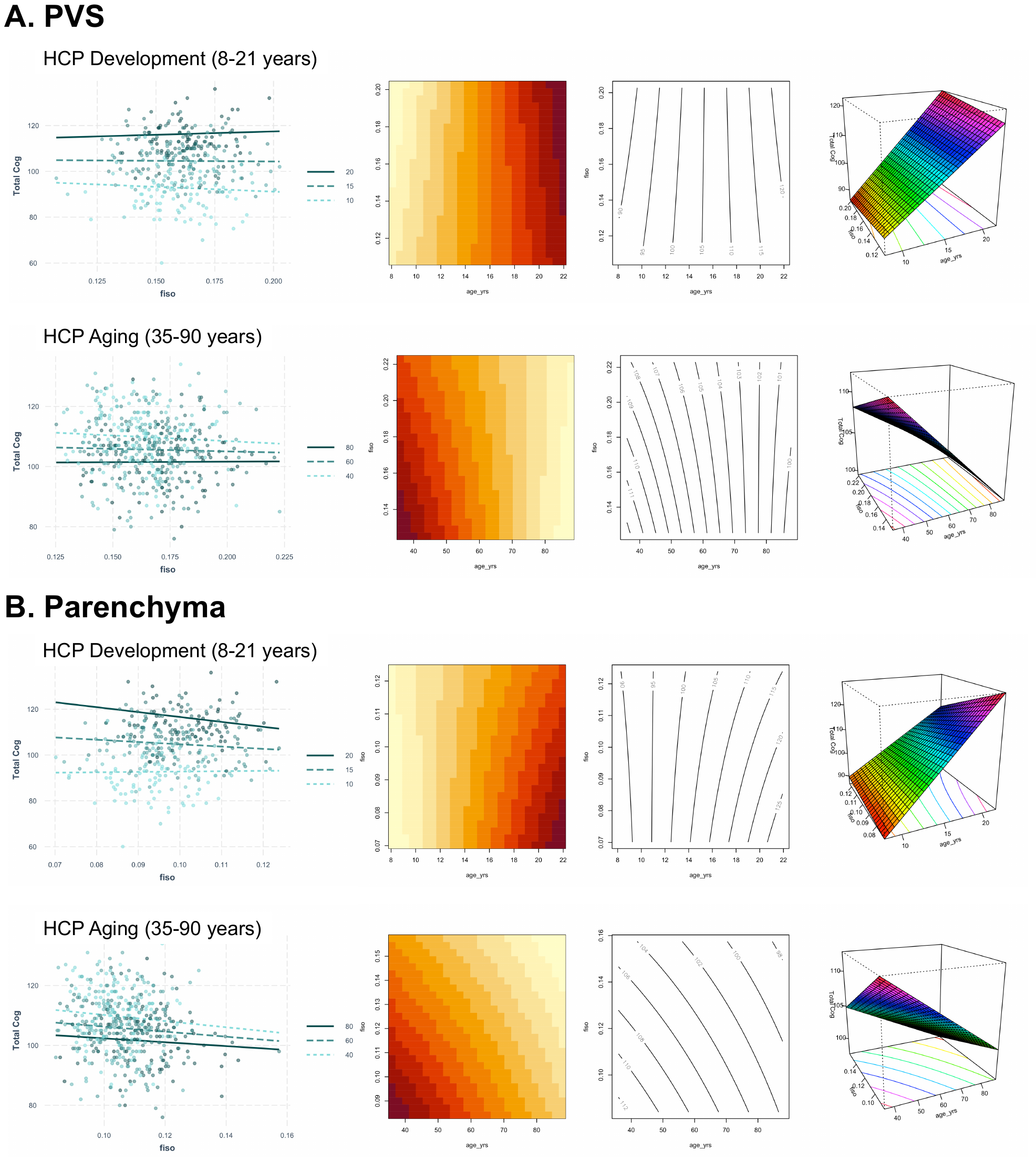

**Supplementary Figure 5**. The relationship between free water diffusion and cognition differs across ages. The relationship between fiso and the NIH total cognition composite score (Total Cog) is shown for the (A) PVS and (B) parenchyma. No significant age*fiso interaction or main effect of fiso was observed on cognition (Left) Individual subjects are shown as points color-coded by age, and representative regression lines are shown for the youngest, middle and oldest ages (from light to dark) for each cohort. (Right) Visualization of the three-dimensional nature of the continuous-by-continuous interactions using matrix plots with contour lines and perspective plots. The contour lines of the matric plots describe the joint distribution of fiso and age, where the area of each binned rectangle in the plot is proportional the frequency of the corresponding combination of variables. The parallel contour lines for both cohorts demonstrates a simple additive relationship. The perspective plots provides a 3D surface-based representation where the height of the surface at any point (x,y = age, fiso) represents the predicted value of the outcome variable (z = cognition). Similarly, the planar surface observed in both cohorts indicates a simple additive relationship.
